# pH-responsive nano-vaccines as antigen and adjuvant carrier that improve vaccine immunogenicity

**DOI:** 10.1101/617324

**Authors:** Daniel Freeman, John Smith

## Abstract

Here, we report a novel method to establish an effective antigen and immunoagonist co-vector to solve the problems of low antigenic endocytosis efficiency, low immunological activity and easy degradation of antigen by antigen presenting cells. Mesoporous silica was selected as the nucleus. After loading the model antigen chicken egg albumin (OVA), the metal organic framework (MOF) formed by Eu ^3+^ and guanine mononucleotide (GMP) was coated on the mesoporous silicon surface. The immunostimulant CpG nucleic acid is adsorbed on the surface to construct a co-carrier system of pH-sensitive antigen and immunostimulant. The antigen loading rate of mesoporous silicon was 20%, and the protein release amount reached 55% after incubation for 24 h under acidic conditions. Transmission electron microscopy showed that the conjugated polymer was uniformly coated on the surface of the material; It was found that the adsorption capacity of the carrier for CpG nucleic acid was 8 ×10^−6^ mol per gram of carrier−adsorbing nucleic acid; MTT results showed that the vector had low toxicity.

## Introduction

The development of vaccines has gone through traditional attenuated or inactivated pathogen microorganisms to subunit vaccines. However, the application of these highly safe protein vaccines is greatly limited without prior in vitro treatment: they are not efficiently taken up by antigen-presenting cells (APCs) and are therefore unable to effectively elicit an immune response. In protein-based immunotherapy, the strategy of nanoparticle-coated antigens has been reported [1□3]. However, most studies have focused on antigen delivery, and few studies have focused on improving the ability of nanocarriers by altering the efficacy of the no-regulator. Co-loading an antigen with an immunological adjuvant or an immunostimulant in the same nanoparticle can increase the body’s immune response. Based on this, a number of research groups simultaneously co-loaded immunostimulators such as unmethylated CpG nucleic acid [4□5] and PAM3CSK4 lipoprotein [6] with antigen to improve the immune response of the vaccine to the body. Although certain immune effects can be achieved, some practical problems limit their clinical application: low antigen coating efficiency, the use of organic solvents affect antigen stability, complex preparation processes and antigen release mechanisms are not clear. The pH-responsive antigen and immunoagonist CpG nucleic acid co-vectors are capable of greatly promoting the T helper cell type 1 immune response through antigen cross-presentation, enhancing cytotoxic T lymphocyte activity (CLT), for anti-intracellular infection and Tumors have an important role [7□10]. This experiment intends to construct a pH-responsive antigen and immunoagonist co-vector system to solve many difficulties faced by traditional vaccine vectors, and provide a theoretical basis for the clinical application of nano-vaccines.

## Materials and Method

### Experimental reagents

Cetyl ammonium bromide (CTAB), tetraethyl orthosilicate (TEOS) and 3 aminopropyltriethoxysilane (APTES) were purchased from Sigma, USA. Chicken ovalbumin (OVA), guanine mononucleotide (GMP) and EuCl 3 were purchased from Shanghai Aladdin Biotech Co., Ltd. The 5□ carboxytetramethylrhodamine-labeled CpG oligonucleotide sequence (5’□TCCATGACGTTCCTGACGTT□3’) was synthesized by Shanghai Biotech Co., Ltd. Other reagents were purchased from Tianjin Komi Chemical Reagent Co., Ltd.

### Experimental Instruments

Transmission Electron Microscopy (TEM): F20S□TWIN, FEI, USA. High-speed refrigerated centrifuge: 3□30K, Sigma, USA. Vacuum freeze dryer: FD8□3, US SIM company. IKA Magnetic Stirrer (Germany); AR3130 Electronic Balance (Shanghai). L□IS□RDV1 type constant temperature oscillator (Shanghai). UV-visible spectrometer: UV□2700, Shimadzu Corporation, Japan. Fluorescence spectrometer: F7000, Hitachi, Japan.

### Experimental method

Preparation of mesoporous silicon Weigh cetyl ammonium bromide (CTAB) 0.96 g, weigh triethanolamine 0. 65 mL, stir at 80 °C for 1 h in oil bath Then, add 7. 8 mL of tetraethyl orthosilicate (TEOS), stir for 2 h, then filter by suction, wash once with absolute ethanol and water, and dry at 100 °C for 2 h at 600 °C. Baked for 6 h, the sample was reserved. Synthesis of aminated mesoporous silicon Weigh 20 mg of MSN, add 20 mL of anhydrous toluene as solvent, dissolve in ultrasonic state, measure and add 400 µL of APTES, install the condensing unit at 110 °C. After stirring for 24 h under oil bath conditions, the sample was centrifuged, washed with ethanol, centrifuged once again, placed in a dry box and dried to a powder state, and the sample was set aside. 1. 3. 3 Antigen OVA load Weigh OVA 20 mg, dissolved in 1 mL water, the OVA mother solution concentration is 20 mg / mL, weigh 10 mg aminated MSN, dissolved in 1 mL water, the concentration is 10 mg / mL, take 3 parts of 100 µL of aminated MSN solution, add 100, 200, 400 µL of prepared OVA mother solution, add all three distilled water to make each solution 1 mL reaction system, stir reaction 24 h. The absorbance of the stock solution and the supernatant was measured by an ultraviolet spectrophotometer to calculate the antigen loading. MOF material coated mesoporous silicon material 6 mg of OVA-loaded aminated SiO2 was weighed into a certain amount of water under ultrasonic conditions until a good dispersion state. The prepared GMP and EuCl 3 solutions were separately added under stirring, and after centrifugation for 10 minutes at a temperature of 10 000 r/min, the mixture was washed once with water and centrifuged again. The coating effect was observed by transmission electron microscopy. Antigen release behavior study The OVA-loaded MSN-NH2@ MOF was dissolved in 100 µL of deionized water and divided into two equal parts: (1) Add 3 mL of PBS with pH= 7. 4; (2) Add 3 mL pH = 5 PBS. Samples were taken under stirring, and 300 µL of the sample was taken each time. After centrifugation, the supernatant was stored, and the precipitate was dissolved in 300 µL of the corresponding pH PBS, and added to the original solution. The sampling time was 3 h, 6 h, 9 h, 12 h, and 24 h. In the reference manual, the concentration of protein in the supernatant was determined by BCA (bicinchonininc acid) protein quantification. 1. 3. 6 CpG Nucleic Acid Adsorption A fluorescently labeled CpG stock solution with a concentration of 0. 2 mol / L was prepared. Add 10 µL to 1 mL of water and dilute it 5 times. Add 0. 1 mg of MSN □NH2@ MOF to react for 24 h under stirring. The supernatant was centrifuged at 10 000 r/min and at a normal temperature for 5 min, and the fluorescence intensity of the CpG nucleic acid in the mother liquid and the supernatant was measured by a fluorescence spectrophotometer to calculate the nucleic acid loading rate. Cytotoxicity study of 1.3. 7 nanocarriers The effect of MSN□NH2@ MOF on the viability of mouse macrophages (RAW264. 7) was examined by MTT assay. The procedure is as follows: Adjust RAW264. 7 Density of 5× 10 5 /well inoculated into 96-well plate, add MSN□NH2@ MOF with final concentration of 25, 50, 100, 200, 400 and 800 µg / mL, PBS as blank Group, each group is set to 6 parallel. After 48 h, add 10 µL of MTT (5 mg / mL) and incubate for another 4 h at 37 °C. The supernatant was discarded, 100 µL of DMSO was added to each well, and the formazan produced by the reaction was dissolved, and the OD value at 570 nm was measured with a microplate reader.

## Results

### Mesoporous silicon in-situ coating MOF material

In order to detect the morphology and coating characteristics of mesoporous silica coated MOF nanoparticles, the synthesized materials were observed by transmission electron microscopy, as shown in Fig. 1. Figure 1a and Figure 1c show the TEM image before MSN coating, which shows that the mesoporous silicon is monodisperse and has a good mesoporous structure with a particle size of about 100 nm. Figure 1b and Figure 1d show the TEM of the mesoporous silicon coated MOF material. It can be seen that the mesoporous silicon pores are uniformly and fully coated by the MOF material, and the coating effect is good.

**Figure 1.**
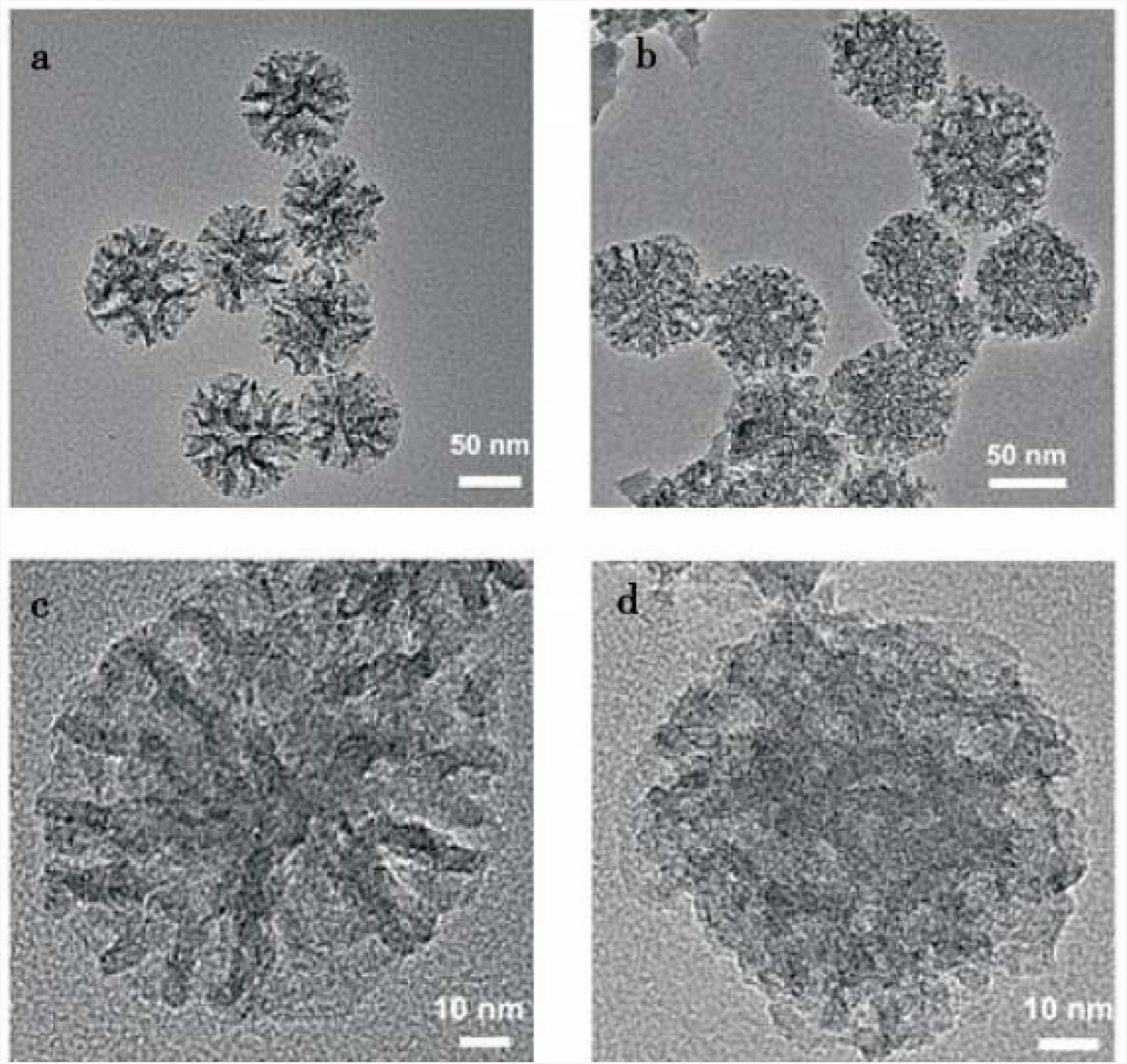
TEM images of pH-responsive nano-vaccines.

### Mesoporous silicon loading OVA

Figure 2 shows the MSN-NH2 drug loading. The abscissa is the amount of antigen (mg) per ml of solution, and the ordinate is the loading rate of the protein. It can be seen from the figure that the adsorption rate reaches a maximum of 20% when the OVA concentration is 4 mg / mL.

**Figure 2.**
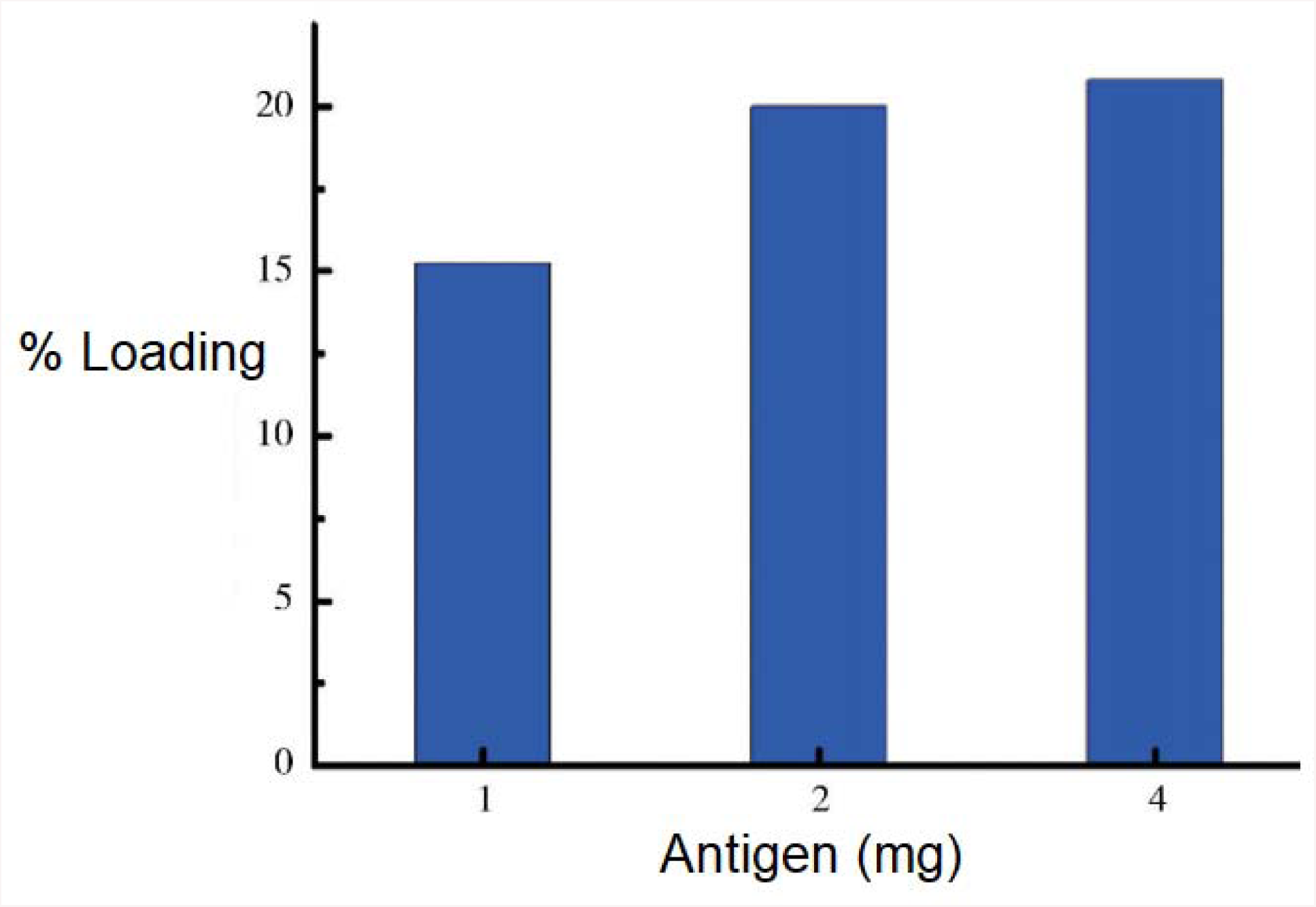
Nano-vaccine Loading capacity.

### MOF material pH-responsive protein release studies

Figure 3 shows the release profile of MSN-NH2@ MOF at different pH states. The concentration of the protein was determined by BCA protein quantitation by sampling at different time points. It can be seen from the figure that the protein release tendency under acidic conditions is more obvious than under neutral conditions. It was demonstrated that MOF released more antigen after dissociation under acidic conditions.

**Figure 3.**
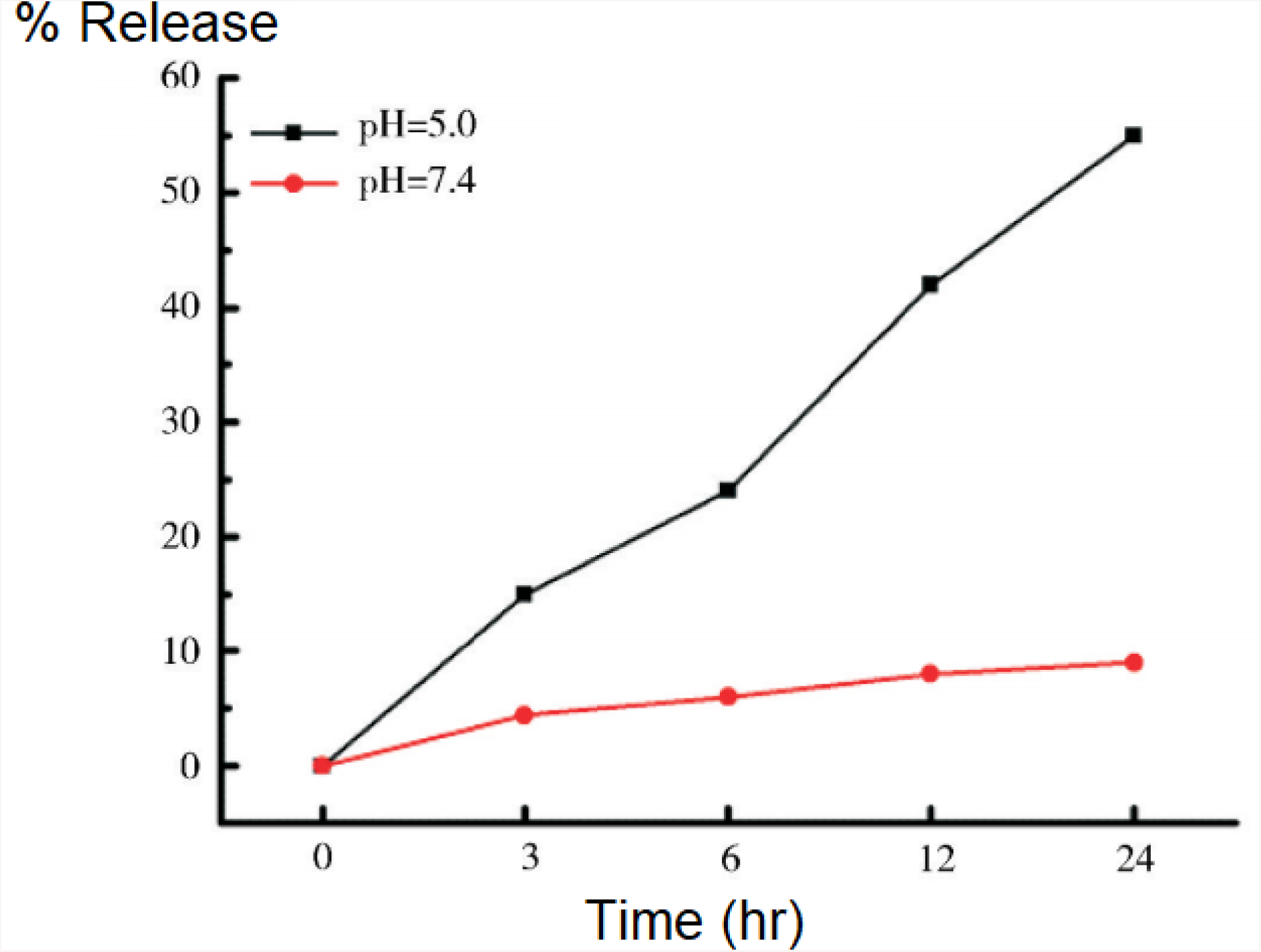
Nano-vaccine loading kinetics.

### CpG immunostimulant adsorption

The adsorption efficiency of MSN-NH2@ MOF on CpG nucleic acid was analyzed by measuring the concentration of Tamra fluorescently labeled CpG in the stock solution and supernatant. Figure 4 shows that as the concentration of nucleic acid increases, the adsorption of nucleic acids by the carrier is gradually enhanced.

**Figure 4.**
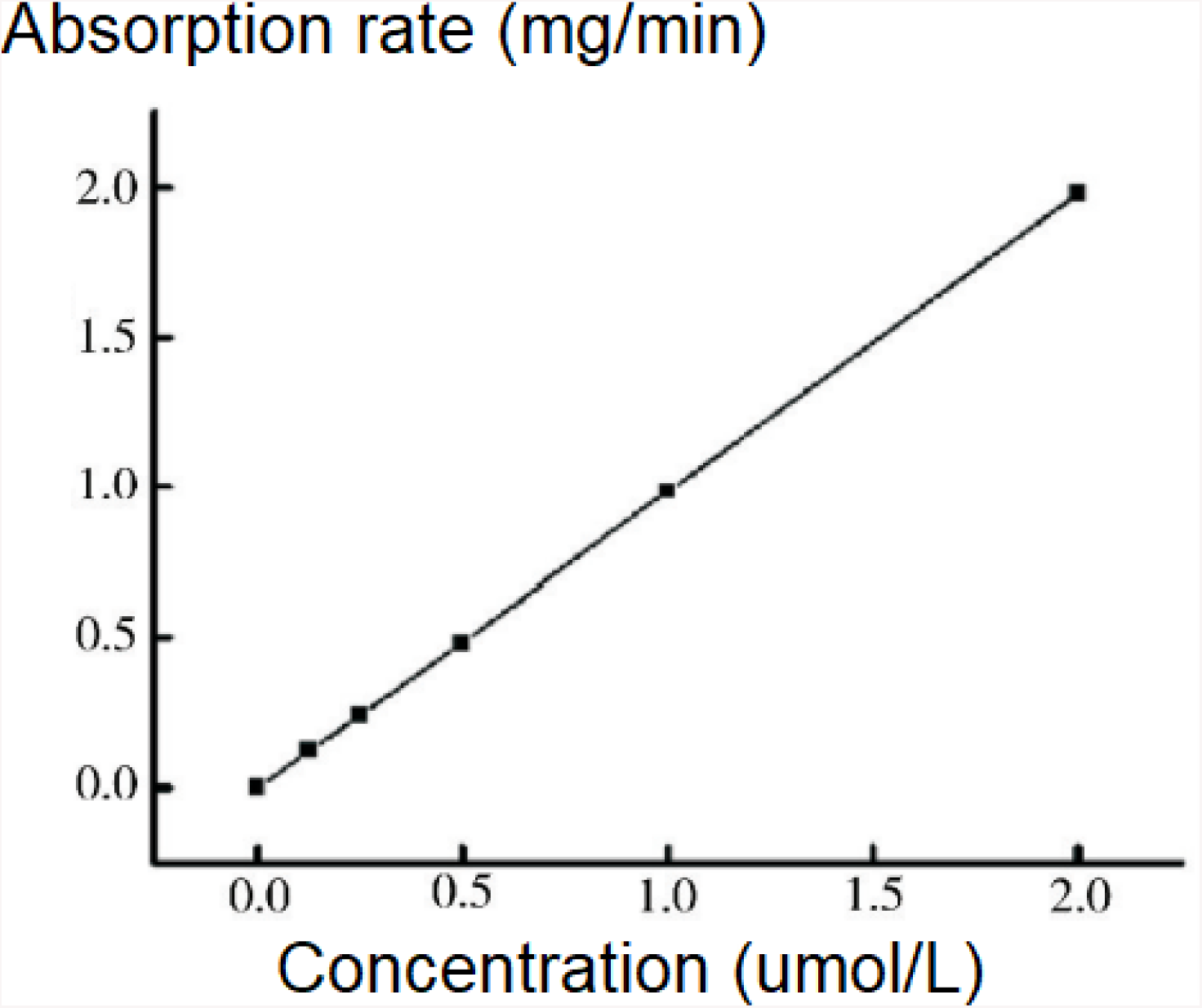
Antigen is absorbed by nano-vaccines in a linear correlation.

### Cytotoxicity study

The effect of MSN-NH2@ MOF on cell viability was evaluated by MTT assay. The MTT results (Fig. 5) showed that the inhibitory effect of the vector on RAW264.7 increased after 48 h of action as the concentration of the vector increased. However, even at high carrier concentrations (800 µg / mL), the cell viability of macrophages is still above 85%. It can be seen that the nanocarrier has good biocompatibility.

**Figure 5.**
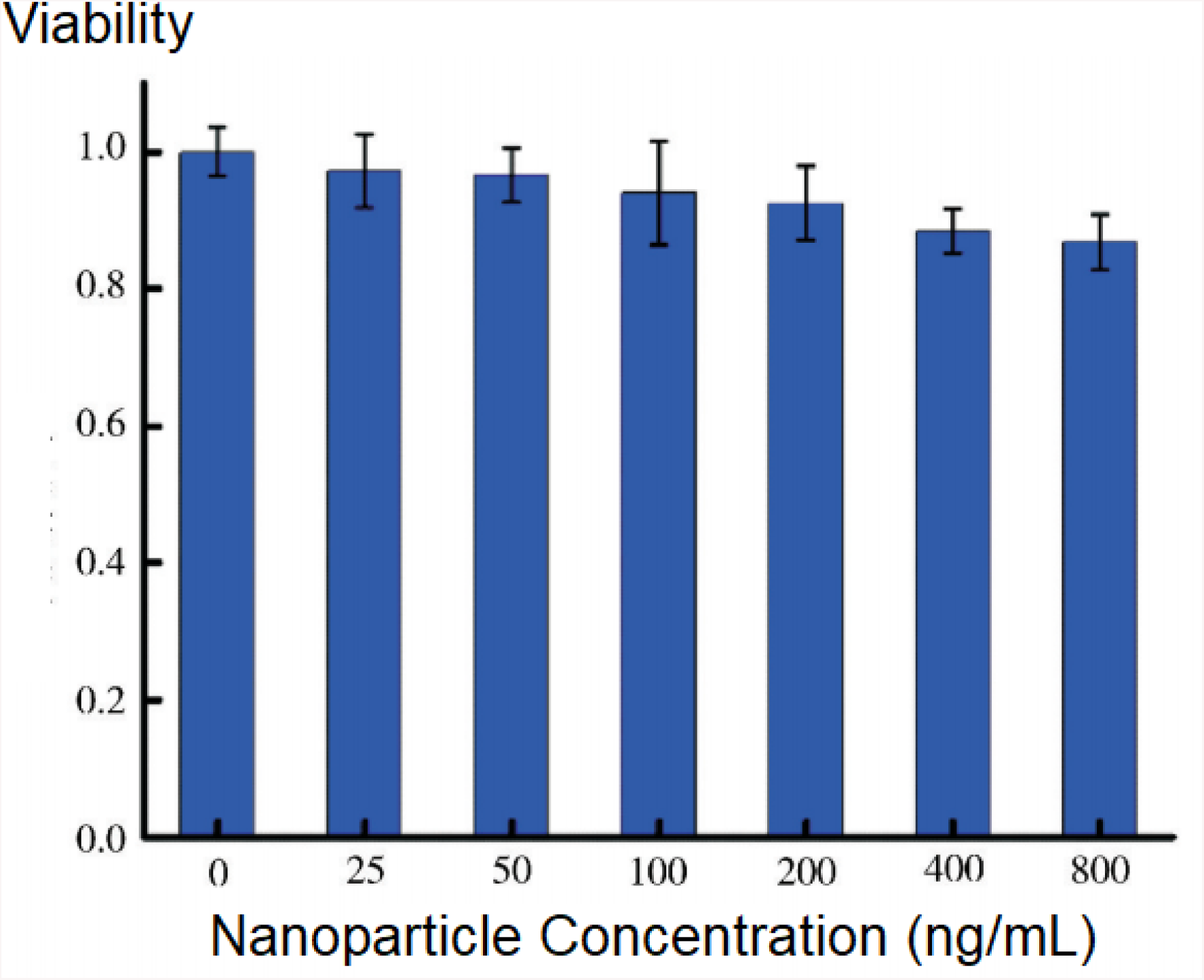
Nanoparticle cytotoxicity assessment.

## Discussion

Infectious diseases have always been considered to be one of the most important hazards that threaten human survival and health and hinder social and economic development. Its prevalence in a society or region is an important indicator of the health of a population in that society or region, and keeping the harm of infectious diseases to a minimum is an important guarantee for sustainable socio-economic development. As the most important means of preventing infectious diseases, vaccines have always received attention. However, traditional inactivated or attenuated pathogen vaccines pose significant risks, and protein-based subunit vaccines developed in recent years have been favored. However, these highly safe protein vaccines are not effectively ingested by antigen presenting cells (APCs) without pretreatment, and are easily degraded in the body, severely limiting their use. Nanocarriers can effectively protect the vaccine from enzymatic degradation by the body, increase the internalization efficiency of APCs, and greatly promote the use of protein-based subunit vaccines^1−14^.

The MSN-loaded subprotein vaccine was used to construct a pH-sensitive antigen and immunostimulant co-vector system to effectively protect the antigen from enzymatic degradation and enhance its immunogenicity. The results show that the carrier designed by the research group can effectively load the antigen, and the MOF material can be uniformly coated in the pores of the MSN to prevent the protein vaccine from being degraded by the protease in the body before entering the antigen presenting cell, and the antigen can be reached in an acidic environment. Effective release. The synthesized materials were analyzed by MTT experiments to show high biocompatibility. In order to further improve the immunostimulatory ability of the constructed nano-vaccine, this experiment adsorbed CpG nucleic acid with immunopromoting function on the surface of MSN-NH2@ MOF, and realized the co-loading of antigen and immunoagonist. The pH-responsive nano-vaccine carrier can not only protect the stability of the antigen, but also improve the release efficiency of the antigen in the cell, and solve the problem that the release mechanism of the nano-vaccine carrier antigen is not clear and the release efficiency is low. The use of immunostimulatory and non-toxic immunostimulants to assist the antigen to enhance the immunogenicity of the vaccine can further improve the body’s immune efficiency. Antigen and adjuvant co-loading systems with pH response have broad clinical application prospects.

## Conclusion

Mesoporous silicon has a high loading rate on antigen, and the conjugated polymer is successfully coated on the surface of mesoporous silicon, which not only protects the antigen, but also adsorbs the immunoagonist nucleic acid on the surface of the material. More importantly, the carrier has the ability to respond to pH response. Future efforts will focus on translating this promising results in a clinical setting.

